# Rapid fluctuations in functional connectivity of cortical networks encode spontaneous

**DOI:** 10.1101/2021.08.15.456390

**Authors:** Hadas Benisty, Daniel Barson, Andrew H. Moberly, Sweyta Lohani, Ronald R. Coifman, Gal Mishne, Michael C. Crair, Jessica A. Cardin, Michael J. Higley

## Abstract

Experimental work across a variety of species has demonstrated that spontaneously generated behaviors are robustly coupled to variation in neural activity within the cerebral cortex. Indeed, functional magnetic resonance imaging (fMRI) data suggest that functional connectivity in cortical networks varies across distinct behavioral states, providing for the dynamic reorganization of patterned activity. However, these studies generally lack the temporal resolution to establish links between cortical signals and the continuously varying fluctuations in spontaneous behavior typically observed in awake animals. Here, we took advantage of recent developments in wide-field, mesoscopic calcium imaging to monitor neural activity across the neocortex of awake mice. We develop a novel “graph of graphs” approach to quantify rapidly time-varying functional connectivity and show that spontaneous behaviors are represented by fast changes in both the activity and correlational structure of cortical network activity. Combining mesoscopic imaging with simultaneous cellular resolution 2-photon microscopy also demonstrated that the correlations among neighboring neurons and between local and large-scale networks also encodes behavior. Finally, the dynamic functional connectivity of mesoscale signals revealed subnetworks that are not predicted by traditional anatomical atlas-based parcellation of the cortex. These results provide new insight into how behavioral information is represented across the mammalian neocortex and demonstrate an analytical framework for investigating time-varying functional connectivity in neural networks.

## Main Text

Cognitive functions such as perception and attention require the dynamic activity of neuronal networks defined by synaptic connectivity over local and long-range spatial scales ^1–3^. Moreover, animals cycle through behavioral states categorized by a variety of physical markers, including pupil dilation, facial movement, and locomotion, that are associated with changes in cognitive performance ^4–9^. Such variation in motor behaviors and arousal are themselves strongly coupled to fluctuations in neural activity, and several recent studies have demonstrated clear modulation of spontaneous and sensory-evoked firing rates associated with transitions between behavioral states ^10–14^. Work in both human and non-human subjects suggests that the spatiotemporal correlations between neural signals in large-scale networks spanning multiple brain regions also vary with state transitions ^20^. These correlations are often viewed as functional connectivity between nodes in a network, reflecting either true structural (synaptic) connections or common inputs ^21^. From these results, behavioral state transitions might be viewed as categorical shifts between stable network configurations optimized for contextually-relevant cognitive functions.

Most analyses of state-dependent functional connectivity in cortical networks rely on binning data within identified epochs (e.g., sleep versus wakefulness or quiescence versus arousal), ignoring the potential for rapid, continuous variation in both behavioral state and network connectivity. As neurons can be exquisitely sensitive to patterned or synchronized input ^22^, it seems reasonable to hypothesize that rapid changes in network correlations are closely linked to the integrative function of single cells. Furthermore, the time-varying correlation between two neuronal signals is a non-linear, second-order function of those signals, that expresses the dynamics of connectivity. Thus, mathematically, activity and dynamic correlation might differentially represent behaviorally relevant information. Nevertheless, the relative contributions of dynamic neural activity versus dynamic inter-node correlations to decoding behavior is unknown.

To explore this question, we used widefield mesoscopic and cellular resolution 2-photon calcium (Ca2+) imaging, both independently and simultaneously, to monitor neural activity in the awake, head-fixed mouse. We developed a novel methodological strategy to view the dynamic correlations between individual cortical parcels and/or neurons as a “graph of graphs”, allowing us to extract their time-varying latent variables through diffusion embedding ^23^. We then asked how accurately these signals could be used to decode spontaneous fluctuations in behavior measured by pupil dilation, facial movements, and locomotion. Our results show that time-varying network correlations carry significant information about behavioral metrics that can outperform analyses based on the dynamics of activity alone. In addition, the dynamic multimodal correlations between local cellular and large-scale networks are also significantly predictive of behavior. We then show that both cortical parcels and single neurons are dynamically correlated to one of two broad mesoscale networks that are distinct from traditional anatomical boundaries. Overall, these findings demonstrate that rapid fluctuations in functional connectivity across spatial scales provide a robust representation of spontaneous behavior.

## Results

### Mesoscopic imaging of spontaneous behavior and cortical network dynamics

We carried out widefield, mesoscopic calcium imaging ^3^ in awake, head-fixed mice expressing the red fluorescent indicator jRCaMP1b ^24^ (Figure 1a). Indicator expression was mediated by neonatal injection of wild-type mice with AAV9-Syn-jRCaMP1b (see Methods) ^25,26^. We simultaneously monitored cortical activity and spontaneous behavioral metrics including pupil diameter, facial movements, and locomotion (Figure 1b-f, see Methods) ^12,13^. Previous studies have often relied on categorical definitions of behavioral state, averaging within epochs according to thresholding of motor signals ^9,13^. However, analysis of our data indicates that, rather than falling into discrete clusters, these metrics are continuously distributed across a range of rapidly varying values (Calinski-Harabasz Index values vs. # of clusters for K-means clustering using 2-6 clusters, p=0.99, ANOVA, Figure 1b-c). We also find that these signals are only modestly correlated with each other (Figure 1d, Supplemental Figure S1), suggesting that they represent underlying latent variables corresponding to distinct behavioral dynamics.

**Figure 1.**
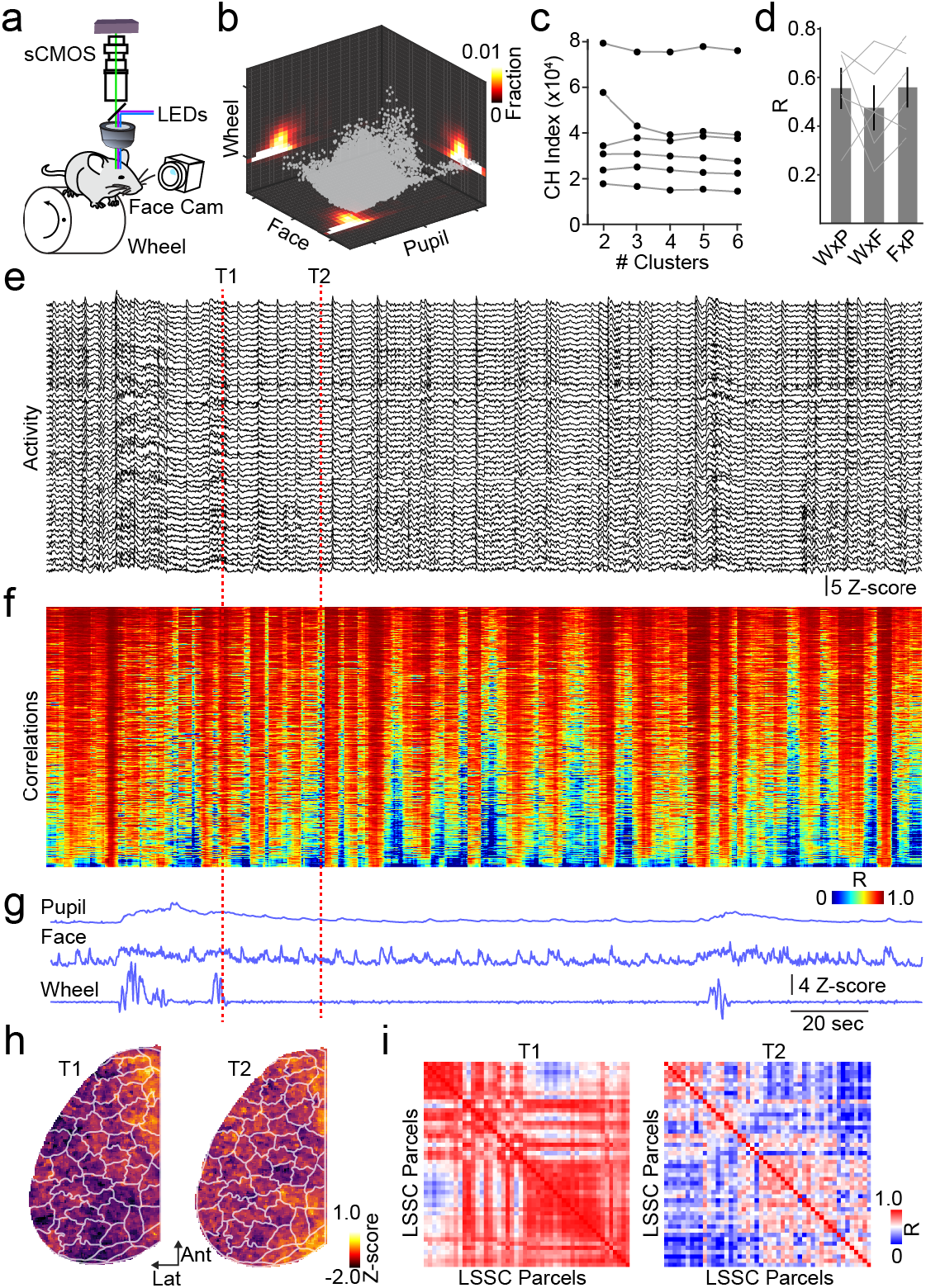
Mesoscopic imaging of cortical activity and functional connectivity. **a**, Schematic illustrating the setup for simultaneous behavioral monitoring and mesoscopic calcium imaging. **b**, Scatter plot illustrating the distribution of Z-scored behavioral metric values (locomotion, facial movement, and pupil size) collapsed over time and across all subjects (n=6 mice). **c**, Population data showing Calinski-Harabasz index values for K-means clustering of behavioral metrics for all subjects. **d**, Population data showing average (±SEM) Pearson’s R^2^ values for the relationships between wheel (W), pupil (P), and facial movements (F) for all subjects. e, Example time series from one animal showing cortical activity across the cortex. Each trace corresponds to one LSSC-based parcel. **f**, Heat map illustrating the time-series of pairwise correlations between each parcel from (e). Data are sorted by increasing standard deviation. **g**, Time series of behavioral metrics corresponding to the data shown in (e) and (f). h, Example LSSC-based functional parcellation of the neocortex for the data shown above. Left and right images are for the timepoints indicated by vertical red lines. *i*, Example pairwise correlation matrices for the data in (e) at the time points indicated.

After normalization and hemodynamic correction of imaging data (see Methods) ^14,27^, we segmented the cortex into functional parcels using a graph theory-based approach that relies on spatiotemporal co-activity between pixels (LSSC, Figure 1e-h, Supplemental Figure S1) ^28^. This approach yields significantly smaller reconstruction errors for a similar number of parcels in comparison to the Allen Institute CCFv3 anatomy-based atlas ^29^ (Supplemental Figure S1). Moreover, LSSC performs similarly to principal component analysis but with the distinct advantage of spatial interpretability, as LSSC generates discrete, disjointed parcels that can provide straightforward links between function and structure (Supplemental Figure S1). We next extracted ***x**_t_*, the time-varying fluctuations in the fluorescence signal associated with each parcel (Figure 1e, see Methods). As expected, variation in activity appeared to be coupled to changes in behavioral metrics over rapid (sub-second) time scales (Figure 1e-h). We then calculated the time-varying, pair-wise correlations *Ĉ_t_* between LSSC parcels using a sliding 3-second window (0.1 second step-size, Figure 1f-i, see Methods). On average, correlations across the cortex were high (r=0.6±0.03, n=6 mice), but their moment-to-moment values also appeared to co-vary with rapid behavioral changes (Figure 1f-i).

### Dynamic functional connectivity encodes spontaneous behavior

We then considered the assumption that behavior can be represented as a nonlinear function of multi-dimensional neural activity beginning with a model of behavior, *b*(*t*), and derived an approximation using the first two terms of a standard Taylor expansion (see Methods):

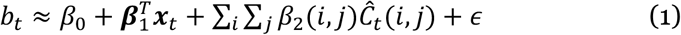

Here, the first-order term ***x**_t_* is an *N*-dimensional vector corresponding to the time-varying neural activity across *N* cortical parcels at time *t*. Indeed, representing behavior as a linear combination of time-varying neural signals is a common approach ^9,30–33^. The second-order term *Ĉ_t_*(*i,j*) corresponds to the time-varying pairwise sample correlations between parcels *i* and *j*. We hypothesized that modelling behavior by combining both a linear term in activity ***x**_t_* and correlations *Ĉ_t_*, which are a nonlinear second-order function of ***x**_t_*, would significantly improve decoding accuracy, suggesting distinct roles for variations in both signal activity and functional connectivity in cortical function.

In an initial effort, fitting a linear ridge regression model for behavioral dynamics whose predictors are time-varying cortical activity and functional connectivity led to poor predictive power (Supplemental Figure 2) due to over-fitting caused by the high-dimensionality of pairwise correlations *Ĉ_t_*(~10^3^ pairs per animal). Therefore, we developed a novel strategy to extract a lower dimensional representation, *ϕ_t_*, capturing the intrinsic dynamics of the correlational signals using Riemannian geometry and diffusion embedding ^23^. Each correlation matrix over a short temporal window (e.g., 3 seconds) can be viewed as a graph whose *N* nodes are cortical parcels connected by weighted edges equal to the instantaneous pairwise correlation coefficients between parcels. Sliding the window over time (Δt=0.1 seconds) produces a series of correlation matrices, which can also be viewed as a time-varying graph (Figure 2a). We then built a “graph of graphs”, where each node is now a time-point represented by the correlation matrix at that time (see Methods). A distance measure between correlation matrices is necessary to set the edge weights of this temporal graph. Since correlation matrices lie on a non-Euclidean Riemannian manifold called the Semi-Positive Definite (SPD) cone, Euclidean distances do not properly represent similarity in this space (Figure 2b). Therefore, we used Riemannian geometry to calculate pairwise geodesic distances between correlation matrices. We applied diffusion embedding to the graph of graphs and extracted the low dimensional representation, *ϕ_t_*, of the temporal dynamics of functional connectivity (Figure 2c).

**Figure 2.**
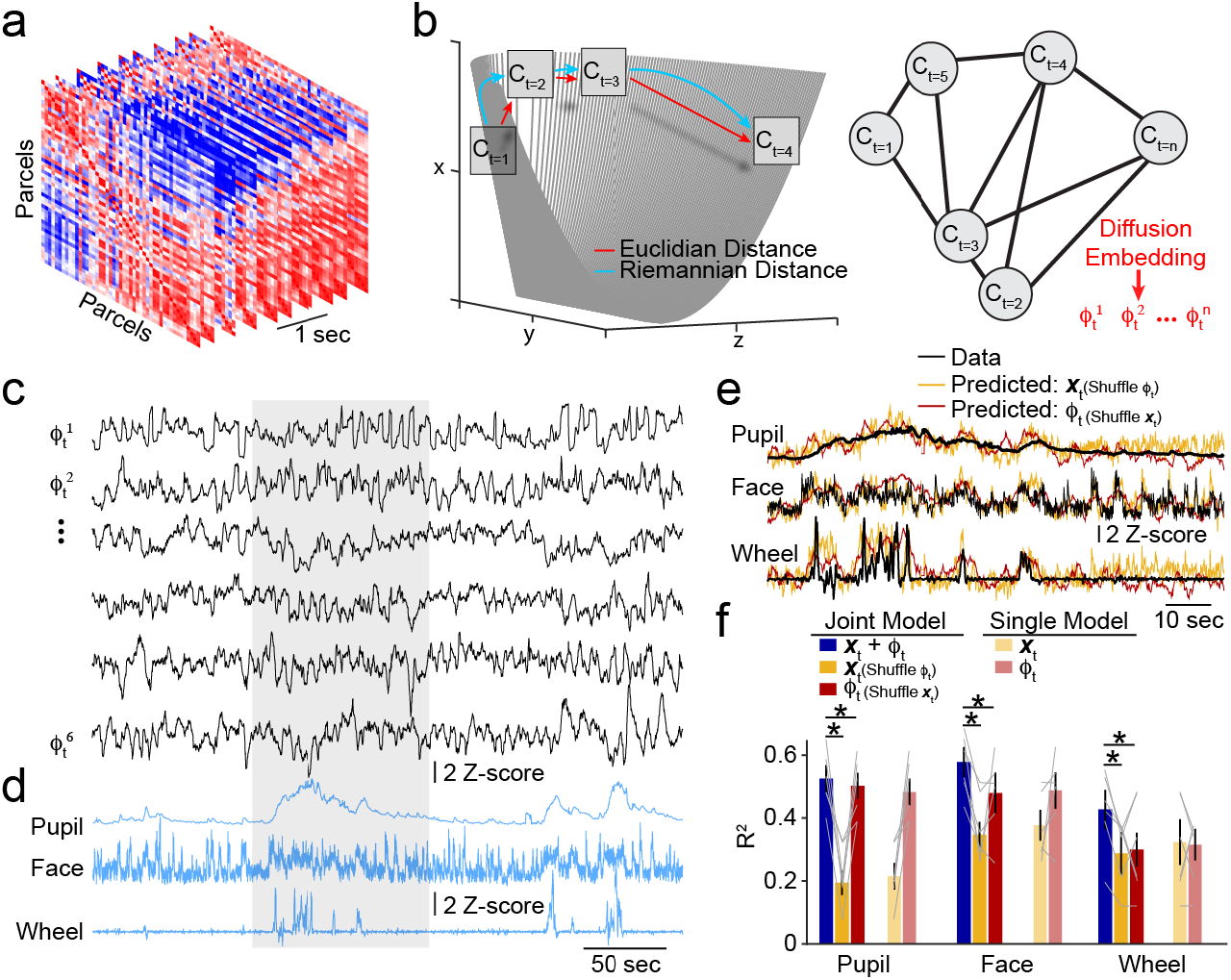
Dynamic functional connectivity encodes rapid behavioral variations. **a**, Example sequential pairwise, parcel-based correlation matrices, derived from a sliding window applied to neural activity across the cortex. **b**, Left, Schematic illustrating the cone-shaped Riemannian manifold (based on 2 × 2 SPD matrices where *x,z* > 0 and *y*^2^<xz, see Methods) used to calculate distances between correlation matrices. The Riemannian measurement reflects geodesic distance that is ignored when using Euclidean distance. Right, Illustration of a “graph of graphs”, whose nodes are SPD matrices and edges are weighted by the length of the geodesic arc along the Riemannian cone, that is used to extract diffusion embedding (*ϕ_t_*) components. **c**, Example diffusion embedding components capturing dynamics of functional connectivity *ϕ_t_*. Simultaneous time series for behavioral metrics are shown below in blue. **d**, Example behavioral data (black traces) showing fluctuations in pupil diameter, facial movement, and locomotion superimposed on predicted behavior estimated using a joint model in which either time-varying activity (red) or embedded correlations (yellow) have been shuffled for the region of data highlighted in (c). **e**, Population data showing average (±SEM) prediction accuracy (R^2^) for modeling behavior variables using a joint model of activity and embedded correlations (blue), joint model with shuffled *ϕ_t_* (yellow), joint model with shuffled activity (red), single predictor model using activity (pale yellow), and single predictor model using *ϕ_t_* (pale red). * indicates p<0.05 (see main text).

We constructed a cross-validated linear regression model combining the cortical activity for all parcels (***x**_t_*, ranging from 48-53 parcels per animal across 6 mice) and the first 20 leading components of the embedded correlations, denoted by 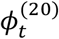 to predict the continuously varying behavioral signals for pupil diameter, facial movement, or locomotion (Figure 2d-e). We found that behavior can be robustly decoded by this joint model (Pupil: R^2^=0.52±0.04; Face: R^2^=0.59±0.04; Wheel: R^2^=0.45±0.06; n=6 mice, Figure 2e-f). As expected, predictive accuracy was significantly impaired when using either raw correlations or Euclidean rather than Riemannian distances for the diffusion embedding (Supplemental Figure S2). Moreover, predictive performance was robust to changes in model parameters, did not improve with the inclusion of more than 20 embedding components, and did not vary appreciably for window lengths of 3-10 seconds (Supplemental Figure S2).

To investigate the relative contributions of activity versus connectivity dynamics in modeling behavior, we recreated a joint model while temporally shuffling one of the two predictors. Shuffling either term significantly impaired prediction accuracy relative to the unshuffled full model (Pupil: R^2^=0.2±0.04, p=0.001 for shuffling 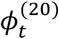, R^2^=0.48±0.04, p=0.02 for shuffling ***x**_t_*; Face: R^2^=0.38±0.05, p=0.002 for shuffling 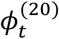, R^2^=0.49±0.06, p=0.006 for shuffling ***x**_t_*; Wheel: R^2^=0.31±0.07, p=0.01 for shuffling 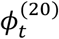, R^2^=0.32±0.05, p=0.003 for shuffling ***x**_t_*; Paired t-test, Figure 2e). Surprisingly, models in which correlational data was preserved were similar or better at decoding behavior than activity-preserved models, reaching significance for variations in pupil diameter (Pupil: p=0.002; Face: p=0.06; Wheel: p=0.37; Paired t-test, Figure 2d-e). To further examine the ability of activity or connectivity signals to independently predict behavioral signals, we generated single predictor models which produced similar results (Figure 2e). Finally, we note that, while the time-averaged activity and pairwise correlations significantly differ for high versus low behavioral state epochs (see Methods), the temporal dynamics of averaged cortical activity (across all parcels) or correlations (across all pairs of parcels) are poorly predictive of the rapid fluctuations in behavior (Supplemental Figure S2). Altogether, these findings indicate that inclusion of rapidly time-varying functional connectivity significantly improves decoding power for modeling of behavioral state, suggesting that cortical network function relies not only on the absolute amount of activity but also on the dynamic coordination of activity across widespread areas.

### Network connectivity does not encode sensory information

Spontaneous cortical activity likely reflects latent signals corresponding to internally generated brain processes. Thus, we asked whether extrinsic sensory information was similarly represented by large-scale networks. We presented the mouse with a series of visual stimuli (drifting sinusoidal gratings, see Methods) and quantified evoked activity using mesoscopic calcium imaging. Contrast-dependent responses were largest in visual areas but were also observed broadly across other cortical regions. However, evoked responses had minimal impact on the correlational structure of activity across the cortex (Supplemental Figure S3). Linear modeling showed that the stimulus could be robustly decoded using activity in visual cortex, with prediction accuracy exhibiting contrast-dependence. However, mesoscopic correlational structure was only very weakly predictive of the stimulus, differing significantly from the activity-based model (F= 57.1, p=5.5e^−10^, ANOVA for combined stimulus contrasts >50%, n=6 mice; Supplemental Figure S3). These results suggest that brief sensory inputs drive large fluctuations in cortical activity without substantial alteration of functional network connectivity, consistent with previous findings that sensory inputs do not interrupt ongoing internal dynamics^12^.

### Dynamic correlations of cellular networks also predict behavior

To determine the generalizability of our approach and also examine encoding by neural correlations at a different spatial scale, we monitored local circuit activity using cellular resolution 2-photon calcium imaging of GCaMP6s-expressing neurons ^34^ in the primary visual cortex (see Methods, Figure 3a-b). As above, we looked at both time-varying activity ***x**_t_* and embedded pair-wise correlations *ϕ_t_* for identified neurons simultaneously with measurements of pupil diameter, facial movement, and locomotion (Figure 3c-f). Unlike large-scale network signals, time-averaged correlations between neurons were broadly distributed around zero (R=−0.001±0.006, n=6 mice).

**Figure 3.**
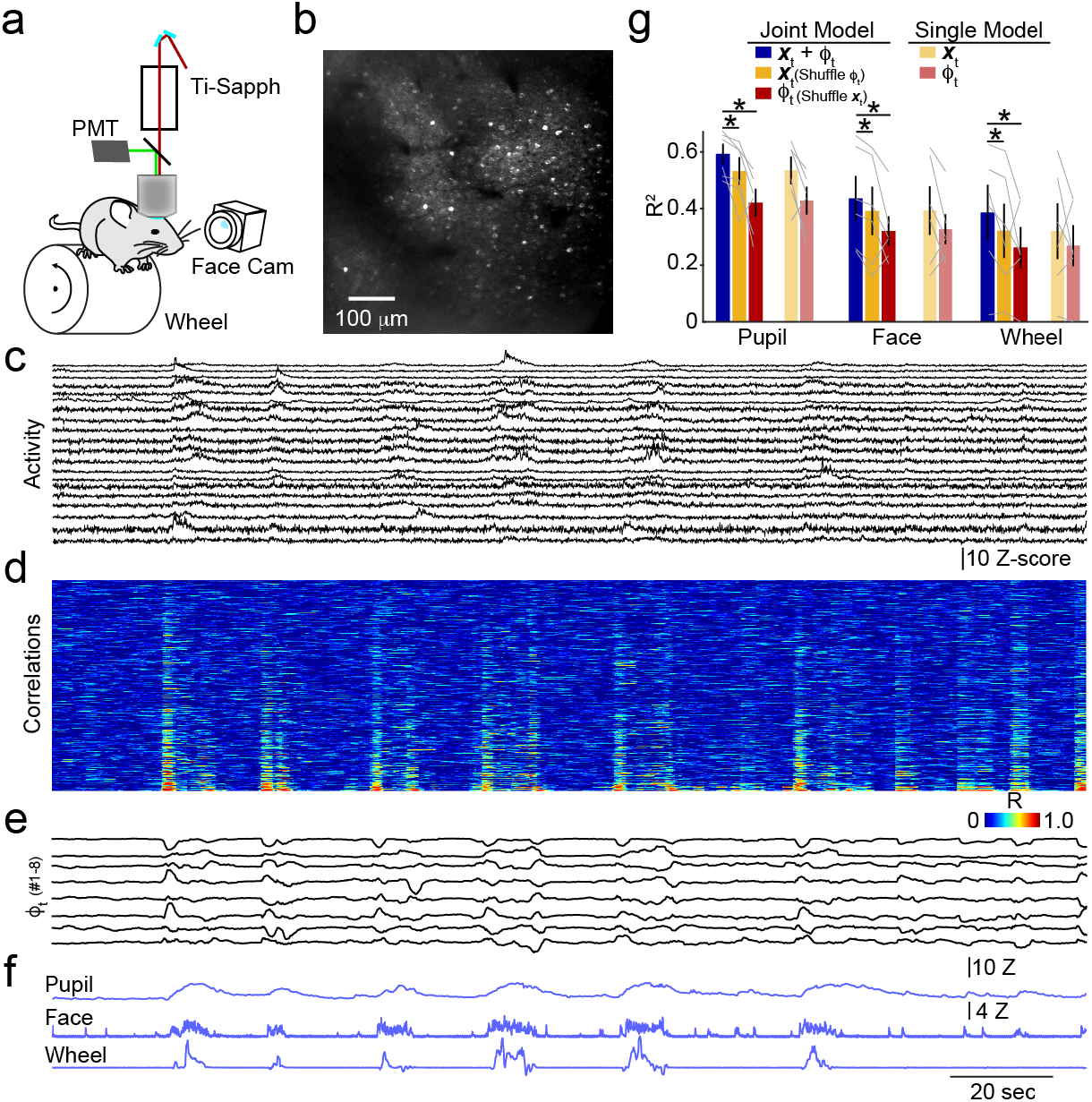
Local circuit dynamics encode spontaneous behavioral variation. **a**, Schematic illustrating the setup for simultaneous behavioral monitoring and 2-photon calcium imaging. **b**, Example field of view showing individual GCaMP6s-expressing neurons in visual cortex. **c**, Example time series showing neuronal activity for all neurons in the field of view. **d**, Heat map illustrating the time-series of pairwise correlations between each neuron from (c). Data are sorted by increasing standard deviation. **e**, Example of the first six diffusion embedding components based on data in (d). **f**, Time-series for behavioral metrics corresponding to data in (c-d). **g**, Population data showing average (±SEM) prediction accuracy (R^2^) for modeling behavior variables using a joint model of activity and embedded correlations (blue), joint model with shuffled *ϕ_t_* (yellow), joint model with shuffled activity (red), single predictor model using activity (pale yellow), and single predictor model using *ϕ_t_* (pale red). * indicates p<0.05 (see main text).

We then generated a cross-validated linear model combining activity and embedded correlation dynamics across cells and attempted to predict rapid fluctuations in behavior. As with mesoscopic imaging, cellular data also robustly predicted behavior (Pupil: R^2^=0.59±0.04; Face: R^2^=0.44±0.08; Wheel: R^2^=0.39±0.1; n=6 mice, Figure 3). Again, modeling performance was poorer using raw correlations and Euclidean distances for embedding but was robust to changes in model parameters (Supplemental Figure S4).

To calculate the relative contributions of activity versus correlations in the joint model, we similarly shuffled one of the two predictors. As above, shuffling either variable significantly impaired prediction accuracy relative to the unshuffled model (Pupil: R^2^=0.53±0.05, p=0.04 for shuffling 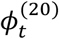, R^2^=0.42±0.05, p=0.0002 for *x_t_*; Face: R^2^=0.39±0.09, p=0.004 for shuffling 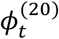, R^2^=0.32±0.05, p=0.03 for shuffling ***x**_t_*; Wheel: R^2^=0.32±0.1, p=0.04 for shuffling 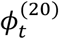, R^2^=0.26±0.07, p=0.02 for shuffling ***x**_t_*; Paired t-test, Figure 3g). Models preserving either the activity or correlational data gave similar accuracy, with activity-based analysis showing modestly better performance for pupil fluctuations (p=0.025 for Pupil, p=0.14 for Face, p=0.15 for Wheel, Paired t-test, Figure 3g). Single-predictor models again produced similar results (Figure 3g). In summary, applying our novel approach for quantifying time-varying correlations in neural data to cellular resolution imaging, we again find that including dynamic functional connectivity significantly enhances prediction accuracy in models linking neural signals to fluctuations in behavioral state.

### Dynamic functional connectivity suggests distinct cortical subnetworks

The improved accuracy of behavioral prediction using embedding of mesoscopic correlation matrices suggests they may reflect underlying principles of structural organization in large-scale cortical networks. We therefore examined the spatial interpretation of *ϕ_t_* by asking how the time-varying correlation for each pair of parcels is represented by the overall embedding. This approach allows us to determine whether the embedding is primarily capturing spatially organized subsets of pairwise correlations. We quantified the goodness-of-fit using 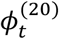 to model the correlation time series between a target parcel and each of the other parcels across the cortex (Figure 4a-b, see Methods). Averaging these goodness-of-fit matrices across all animals (n=6 mice) revealed substantial within-subject spatial heterogeneity that was conserved across different individuals. The embedding primarily represented correlations between each target parcel and one or both of a posterior and anterolateral subdivision of the cortex (Figure 4c, Supplemental Figure S5). This spatial pattern was clearly evident after making a grand average across all parcels and animals (Figure 4d). Intuitively, this result indicates that independent of behavior, the dynamic large-scale correlations of cortical areas are dominated by the interrelationship of each cortical parcel with one or both of these two subnetworks. Surprisingly, this functional organization is highly distinct from regional boundaries defined by traditional, anatomy-based atlases such as the CCFv3 (Figure 4d).

**Figure 4.**
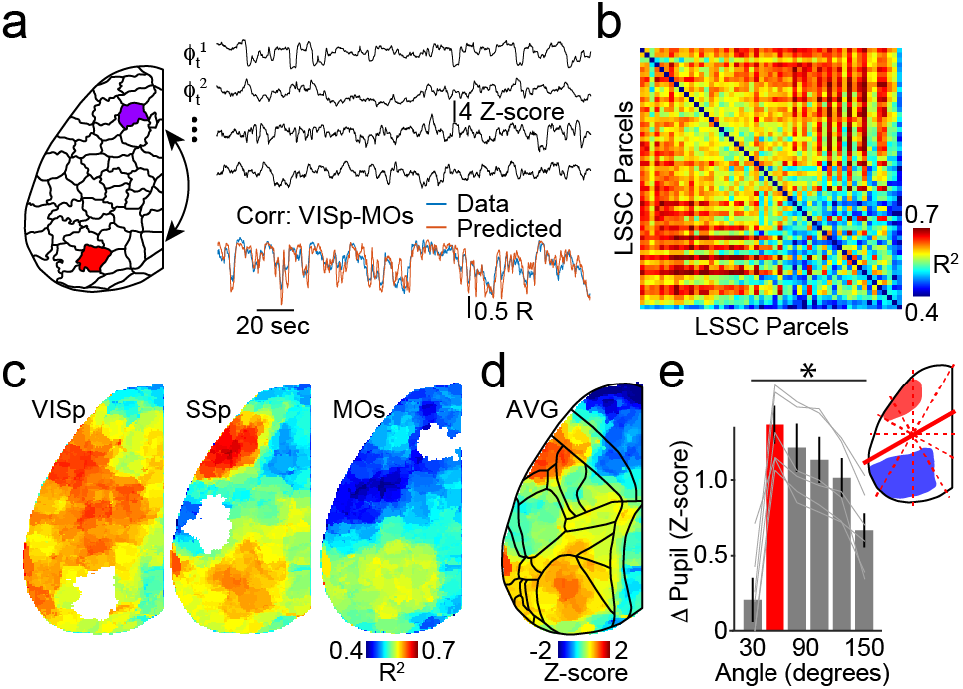
Dynamic functional connectivity reveals distinct cortical subnetworks. **a**, Left illustration of LSSC-based parcellation, highlighting two parcels corresponding approximately to supplemental motor cortex (MOs) and primary visual cortex (VISp) based on CCFv3. Right, example components of correlation embedding for one animal (black), pairwise time-varying correlation between VISp and MOs (blue), and the predicted VISp-MOs correlation based on embedding. **b**, Example matrix from one animal showing the goodness of fit (R^2^) for modeling the time-varying correlations between each pair of parcels using 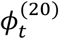. **c**, Average (n=6 mice) maps showing mean R^2^ values for modeling the pairwise correlations of each cortical parcel with the indicated target parcel (shown in white). **d**, Grand average map showing R^2^ values as in (c) collapsed across all animals (n=6) and all cortical parcels. **e**, Population data showing the average (±SEM) difference in pupil size for epochs corresponding to maximal and minimal network segregation versus a line angle bisecting LSSC parcels. 30° (red) corresponds to the division between anterolateral and posterior network (inset). * indicates p<0.05 (see main text).

To further examine whether coordinated activity across this anterolateral/posterior partition corresponds to spontaneous behavioral variation, we calculated a time-varying participation coefficient that measures the functional connectivity of parcels within versus between two groupings ^35^ defined by a line bisecting the cortex (Figure 4e, see Methods). We quantified the difference in average behavioral metrics for time points corresponding to the upper versus lower deciles of the participation coefficients and repeated this analysis for different angles of the bisecting line. Our results showed a significant variation across angles for all behaviors (F=9.5, p=1.6e-5 for pupil; F=5.9, p=0.0006 for Face; F=6.3, p=0.0004 for Wheel; ANOVA, n=6 mice, Figure 4f, Supplemental Figure S5) with the peak value corresponding to the observed boundary of anterolateral and posterior networks. This result supports the conclusion that these subdivisions reflect behaviorally relevant functional cortical architecture.

### Functional connectivity across spatial scales encodes behavior

Our analyses revealed that spontaneous behaviors can be accurately decoded from the temporal dynamics of correlations between neural signals at both local circuit and mesoscopic spatial scales. However single neurons are embedded in large-scale networks by virtue of long-range synaptic connections. Thus, we wondered whether the functional connectivity across these scales was similarly dynamic. To this end, we carried out simultaneous wide-field and cellular 2-photon imaging (Figure 5a-b, see Methods) ^25^. We first analyzed these multimodal data sets separately as described above and quantified the accuracy with which linear models based on time-varying activity ***x**_t_* or embedded correlations 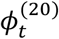 could predict behavior. Activity was predictive for mesoscopic (Pupil: R^2^=0.13±0.03; Face: R^2^=0.14±0.04; Wheel: R^2^=0.12±0.04; n=7 mice) and cellular (Pupil: R^2^=0.31±0.04; Face: R^2^=0.4±0.07; Wheel: R^2^=0.5±0.08) data. Similarly, embedded correlations were also predictive for mesoscopic (Pupil: R^2^=0.35±0.03; Face: R^2^=0.38±0.06; Wheel: R^2^=0.51±0.06) and cellular (Pupil: R^2^=0.14±0.03; Face: R^2^=0.12±0.05; Wheel: R^2^=0.15±0.06) data (Figure 5c). As above, correlations-based prediction accuracy was greater for mesoscopic data (Pupil: p=0.002; Face: p=0.004; Wheel: p=0.0002; paired t-test, n=7 mice) and activity-based prediction accuracy was greater for cellular data (Pupil: p=0.0001; Face: p=0.001; Wheel: p=0.0004; paired t-test, n=7 mice).

**Figure 5.**
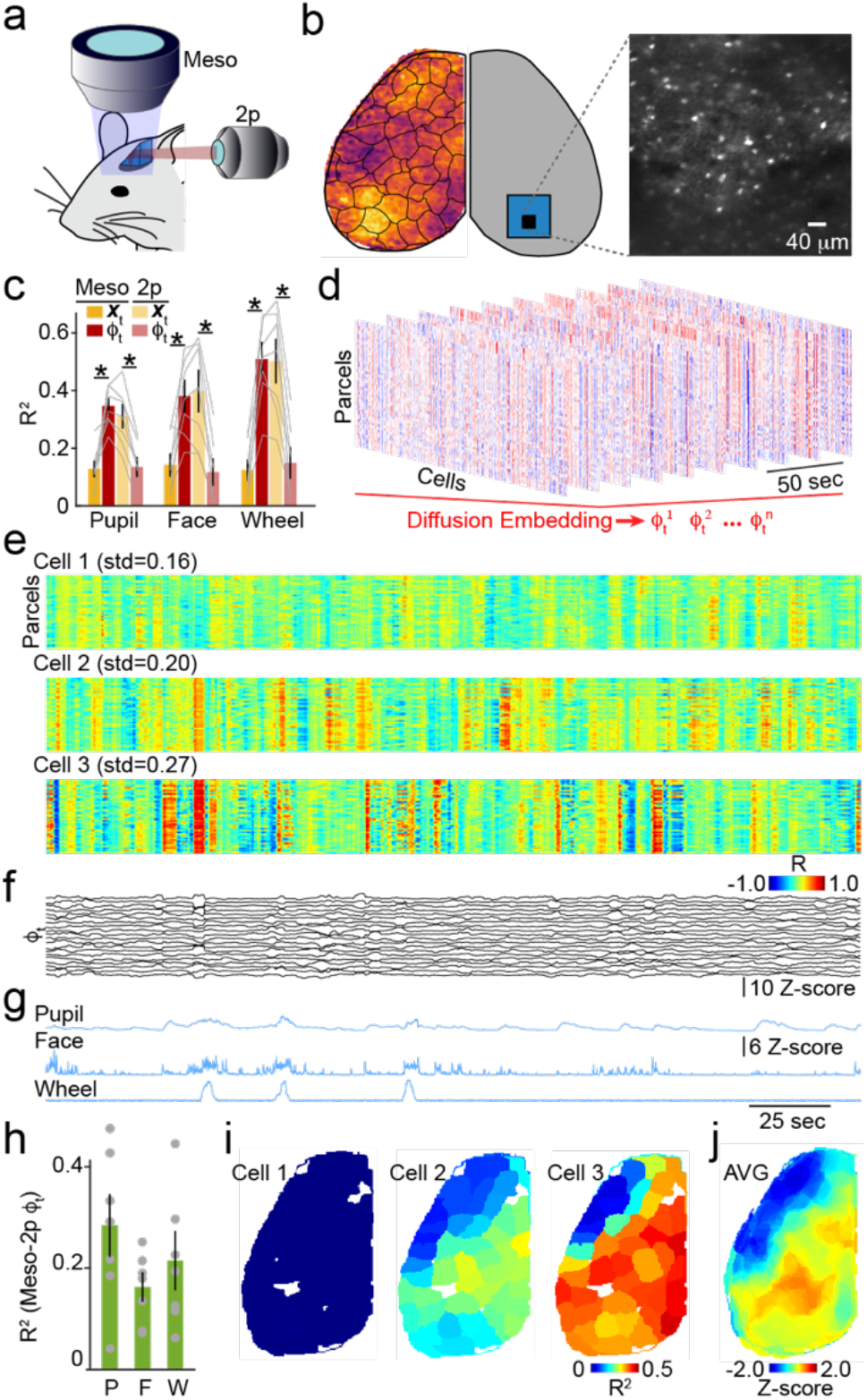
Functional connectivity across spatial scales encodes behavior. **a,** Schematic illustrating the setup for simultaneous mesoscopic and 2-photon imaging. **b,** Left, example mesoscopic imaging frame and schematic of microprism placement in the contralateral hemisphere. Right, example 2-photon imaging frame collected through the prism. **c,** Population data showing average (±SEM) prediction accuracy (R^2^) for modeling behavior variables using either activity (yellow) or *ϕ_t_* (red) for mesoscopic or 2-photon data. **d,** Example sequential multimodal correlation matrices, derived from a sliding window applied to neural activity from mesoscopic (parcels) and 2-photon (cells) imaging, used for diffusion embedding. **e**, Dynamic multimodal correlation time series for three example cells, where each row represents a mesoscopic parcel. The standard deviation of correlation values over time, averaged across all rows is indicated. **f**, Example of the first 20 diffusion embedding components from the same animal (n=243 cells, 47 parcels). **g,** Time series for behavioral metrics corresponding to data in (e-g). **h,** Population data showing average (±SEM) prediction accuracy (R^2^) for modeling behavior variables using *ϕ_t_* derived from the embedding of dual mesoscopic and 2-photon correlations. **i**, Example maps for the cells in (e) showing R^2^ values for modeling the correlation of the cell with each parcel using the overall diffusion embedding. **j**, Grand average map showing R^2^ values as in (i) collapsed across all animals (n=6) and all cells and cortical parcels.

To explore functional connectivity across spatial scales, we developed a strategy to calculate the time-varying correlations between cells and parcels for a sliding 3-second window followed by diffusion embedding (Figure 5d-e, see Methods). This analysis revealed considerable heterogeneity in the degree of multimodal correlation dynamics exhibited by different cells, measured as the standard deviation of correlation fluctuations averaged for a single cell across all brain parcels across the imaging session (Figure 5e, Supplemental Figure S6, see methods). The variation in both the correlations and the embedding components appeared to track with behavioral metrics (Figure 5e-g). Indeed, the embedding of the correlations for the dual imaging data could significantly predict fluctuations in pupil diameter, facial movement, and locomotion (Figure 5h), with results robust across a range of model parameters (Pupil: R^2^=0.29±0.06; Face: R^2^=0.16±0.03; Wheel: R^2^=0.22±0.06; n=7 mice, Supplemental Figure S6).

We explored the spatial interpretation of the dual mesoscopic-cellular embedding by quantifying the accuracy with which the correlations between a single neuron and the mesoscopic parcels are represented by the overall embedding. In general, cells with the most dynamic correlations (largest standard deviation) exhibited the strongest spatial heterogeneity in their modeling accuracy (Figure 5i, Supplemental Figure S6). However, the spatial pattern was generally conserved for all cells and was clearly evident after averaging across the population (Figure 5j), again showing a division of the cortex into anterolateral and posterior subnetworks, with cells in visual cortex being dynamically correlated most strongly with the latter. These results provide an additional independent demonstration that these subdivisions reflect a fundamental organizing principle in the cortex.

## Discussion

Our results show that functional connectivity in cortical networks is highly dynamic, varying on a sub-second time-scale that tracks with continuous metrics of spontaneous behaviors. Including these dynamic correlations in a linear model predicted behavioral fluctuations with high accuracy. This result was true for both large-scale networks monitored with mesoscopic calcium imaging and local networks monitored at cellular resolution with 2-photon microscopy. Moreover, combining these modalities revealed that behavior was also accurately represented by the dynamic correlations between local and cortex-wide networks. The spatial organization of dynamic correlations between either parcels or neurons and cortex-wide activity revealed two distinct subnetworks not predicted from standard anatomical divisions.

The representation of behavioral information by time-varying cortical signals has been a focus of recent studies using diverse approaches to monitor brain activity ^1,2,10,15^. In rodent models, variations in behavioral state or arousal are coupled to changes in firing rate, pairwise correlations, and neuromodulatory signaling ^11–14,36,37^. In particular, arousal- and motor-related variables (e.g., pupil diameter, locomotion, whisking) are represented at the cellular and network scale ^11,12,14^. Several groups have also demonstrated that the spatial patterns of large-scale activity in the neocortex are markedly different when comparing across state ^14,16,20^. For example, spontaneous cortical activity can be decomposed using various methods into repeating spatiotemporal motifs that may correspond to sensory or motor signals ^20^. Similarly in human subjects, shifts in wakefulness correspond to changes in average resting state connectivity ^38^, resting state fluctuations predict somatosensory perception ^4^, and working memory-based task performance corresponds to spatially heterogeneous variation in timescales of patterned activity ^39,40^.

Prior efforts to link cortical activity to spontaneous behaviors have often relied on categorical definitions of state, a strategy that is particularly common for methodologies with limited temporal resolution such as fMRI. However, our data indicate that variations in pupil diameter, facial movement, and locomotion do not appear to cluster into distinct regimes, a result more consistent with continuously and rapidly varying states. Recently, both Stringer et al. ^12^ and Musall et al. ^11^ demonstrated that rapid dynamics of behavior variables could accurately encode neural activity across the cortex, also supported by our present results. Here, we further explored the dynamics of functional connectivity expressed as the correlation between cortical parcels or neurons. Intuitively, time-varying activity and pairwise correlations can be viewed as first- and second-order terms in a Taylor expansion of a function relating behavior to neural signals. Thus, correlations cannot be linearly derived from the underlying activity and represent a potential mechanism to encode an independent component of behavioral dynamics. This conclusion is strongly supported by our results, where shuffling either activity or correlations significantly reduces modeling accuracy for both mesoscopic and 2-photon data.

The temporal scale of the neural and behavioral dynamics is similar to synaptic integration windows for single cells, suggesting the hypothesis that neurons may be sensitive to convergent synaptic input driven by correlated large-scale activity. Our results combining mesoscopic and 2-photon imaging demonstrate that the functional connectivity across these divergent spatial scales is also dynamic and accurately predictive of fluctuations in behavior. Thus, we propose that network activity within and across spatial scales in the neocortex is coordinated as a function of spontaneous behaviors. In the near future, ongoing development of multi-modal approaches, combining fluorescence imaging, fMRI, and electrophysiology ^25,41–43^, will likely drive additional discoveries into the functional organization of brain networks in diverse systems.

Our method for viewing time-varying functional connectivity in cortical networks as a graph of graphs provides an analytical framework for extracting the intrinsic dynamics of short-term correlations and uses Riemannian geometry to correctly evaluate distances between correlation matrices extracted at different time points. These distances are then used to set the weights of a graph-of-graphs, allowing us to extract a low-dimensional representation for the manifold of the correlations and capture their underlying dynamics. Using this approach and including both first-order (activity) and nonlinear second-order (embedded functional connectivity) terms for modeling behavior enabled us to significantly improve decoding power. Our method was generalizable for three different data sets, mesoscopic and 2-photon imaging alone and in combination, and yielded the surprising finding that higher-order statistics (i.e., correlational signals) can produce similar or better predictive accuracy than time-varying changes in activity.

Several distinct strategies have been developed to analyze the spatiotemporal organization of network activity, including singular value decomposition and non-negative matrix factorization ^16,44^. Here, we show that functional parcellation of cortical regions ^28^ followed by embedding of time-varying correlations based on Riemannian geometry, provides a robust means to quantify dynamic functional connectivity that accurately decodes spontaneous fluctuations in behavior. With the increasing interest in analysis of neural manifolds, our results also highlight the necessity of considering the geometry of the manifold on which the data lie to accurately reveal their intrinsic representation. Notably, we did not find similarly strong representation of sensory information in mesoscopic correlations, suggesting that external inputs modulate activity without altering functional connectivity. This finding is similar to recent work suggesting spontaneous behavior and external stimuli are represented in orthogonal dimensions ^12^. Such results may also reflect a lack of behavioral relevance for the stimuli as presented here. We and others have shown that training can modify the sensory and motor representations by single cortical neurons ^9,45–47^. and future studies must determine if development, experience, or learning produce functional reorganization of large-scale networks.

Finally, our results suggest that the cortex can be spatially segmented into two broad subnetworks, an anterolateral and posterior division, a functional division that emerges from analysis of spontaneous activity but also reflects variation in behavioral state metrics. We previously suggested such a division based on correlations between single cell activity and mesoscopic cortical signals ^25^. Interestingly, this organization does not map readily onto standard anatomical segmentation of the cortex, such as the CCFv3 ^29^. We propose that the dynamic modulation and plasticity of synaptic strength may support the translation between such structural and functional views of connectivity in cortical networks, a hypothesis that awaits experimental validation.

## Author Contributions

HB, RRC, GM, JAC, and MJH designed the study. HB, DB, RRC, GM, and MJH developed the analytical approach. HB carried out all analyses. DB, AHM, and SL collected experimental data. HB and MJH wrote the manuscript.

## Acknowledgements

The authors thank members of the Higley and Cardin laboratories for helpful input throughout all stages of this study. We thank Rima Pant for generation of AAV vectors. We thank the GENIE Project for jRCaMP1b plasmids. This work was supported by funding from the NIH (MH099045 and MH121841 to MJH, EY022951 to JAC, MH113852 to MJH and JAC, EY029581 and GM007205 to DB, EY031133 to AHM, EY026878 to the Yale Vision Core, EB026936 to GM and RRC), the NSF (CCF-2217058 to GM), an award from the Yale Kavli Institute of Neuroscience (to MJH and RRC), an award from the Swartz Foundation (to HB), an award from the Simons Foundation SFARI (to MJH), and a BBRF Young Investigator Grant (to SL).

## Data Availability Statement

The full datasets generated and analyzed in this study are available from the corresponding authors on reasonable request.

## Code Availability Statement

Custom written MATLAB scripts used in this study are available on github (https://github.com/cardin-higley-lab).

## Materials and Methods

All animal handling and experiments were performed according to the ethical guidelines of the Institutional Animal Care and Use Committee of the Yale University School of Medicine. Mesoscopic imaging data were collected as part of a previous study ^14^, with experimental details provided below for clarity. Analysis results presented here represent wholly new findings and have not appeared elsewhere.

### Animals

Male and female mice were kept on a 12h light/dark cycle, provided with food and water ad libitum, and housed individually following headpost implants. Imaging experiments were performed during the light phase of the cycle. For mesoscopic imaging, brain-wide expression of jRCaMP1b ^24^ was achieved via postnatal sinus injection as described previously ^25,26^. Briefly, P0-P1 litters were removed from their home cage and placed on a heating pad. Pups were kept on ice for 5 min to induce anesthesia via hypothermia and then maintained on a metal plate surrounded by ice for the duration of the injection. Pups were injected bilaterally with 4 ul of AAV9-hSyn-NES-jRCaMP1b (2.5×10^13 gc/ml, Addgene). Mice also received an injection of AAV9-hSyn-GRABACh3.0 to express the genetically encoded cholinergic sensor GRABACh3.0 ^48^, although these data were not used in the present study. Once the entire litter was injected, pups were returned to their home cage. For two-photon imaging experiments, a similar procedure was used to drive cortex-wide expression of GCaMP6s ^34^. For dual mesoscopic and two-photon imaging experiments, adult (P60-70) mice transgenically expressing GCaMP6s in cortical excitatory neurons (CaMK2a-tTA; tetO-GCaMP6s; VIP-Cre) ^49^ were used. These animals were also injected with AAV driving Cre-dependent GCaMP6s and Cre-dependent tdTomato, though all red fluorescent cells were excluded from the present analysis.

### Surgical procedures

All surgical implant procedures were performed on adult mice (>P50). Mice were anesthetized using 1-2% isoflurane and maintained at 37°C for the duration of the surgery. For mesoscopic imaging, the skin and fascia above the skull were removed from the nasal bone to the posterior of the intraparietal bone and laterally between the temporal muscles. The surface of the skull was thoroughly cleaned with saline and the edges of the incision secured to the skull with Vetbond. A custom titanium headpost was secured to the skull with transparent dental cement (Metabond, Parkell), and a thin layer of dental cement was applied to the entire dorsal surface of the skull. Next, a layer of cyanoacrylate (Maxi-Cure, Bob Smith Industries) was used to cover the skull and left to cure <30 min at room temperature to provide a smooth surface for transcranial imaging. A similar procedure was used to prepare mice for two-photon imaging, with the addition of a dual-layer glass window implanted into a small (<4 mm square) craniotomy placed over the left primary visual cortex. The edges of the window were then sealed to the skull with dental cement. For dual mesoscopic and two-photon imaging, a 2mm glass microprism (Tower Optical) was placed on top of a dual-layer glass window implanted over the right primary visual cortex ^25^.

### Mesoscopic imaging

Widefield mesoscopic calcium imaging was performed using a Zeiss Axiozoom with a 1×, 0.25 NA objective with a 56 mm working distance (Zeiss). Epifluorescent excitation was provided by an LED bank (Spectra X Light Engine, Lumencor) using two output wavelengths: 395/25 (isosbestic for GRAB_ACh3.0_) and 575/25 nm (jRCaMP1b). Emitted light passed through a dual camera image splitter (TwinCam, Cairn Research) then through either a 525/50 (GRAB_ACh3.0_) or 630/75 (jRCaMP1b) emission filter (Chroma) before it reached two sCMOS cameras (Orca-Flash V3, Hamamatsu). Images were acquired at 512×512 resolution after 4× pixel binning, and each channel was acquired at 10 Hz with 20 ms exposure using HCImage software (Hamamatsu).

### Two-photon imaging

Two-photon imaging was performed using a MOM microscope (Sutter Instruments) coupled to a 16×, 0.8 NA objective (Nikon). Excitation was driven by a Titanium-Sapphire Laser (Mai-Tai eHP DeepSee, Spectra-Physics) tuned to 920 nm. Emitted light was collected through a 525/50 filter and a gallium arsenide phosphide photomultiplier tube (Hamamatsu). Images were acquired at 512×512 resolution at 30 Hz using a galvo-resonant scan system controlled by ScanImage software (Vidrio).

### Dual mesoscopic and two-photon imaging

Dual imaging was carried out using a custom microscope combining a Zeiss Axiozoom (as above) and a Sutter MOM (as above), as described previously ^25^. To image through the implanted prism, a long-working distance objective (20×, 0.4 NA, Mitutoyo) was used. Frame acquisitions were interleaved with an overall rate of 9.15 Hz, with each cycle alternating sequentially between a 920nm two-photon acquisition (512×512 resolution), a 395/25nm widefield excitation acquisition, and a 470/20nm widefield excitation acquisition. Widefield data were collected through a 525/50nm filter into a sCMOS camera (Orca Fusion, Hamamatsu) at 576×576 resolution after 45× pixel binning with 20ms exposure.

### Behavioral monitoring

All imaging was performed in awake, behaving mice that were head-fixed so that they could freely run on a cylindrical wheel. A magnetic angle sensor (Digikey) attached to the wheel continuously monitored wheel motion. Mice received at least three wheel-training habituation sessions before imaging to ensure consistent running bouts. During widefield imaging sessions, the face (including the pupil and whiskers) was illuminated with an IR LED bank and imaged with a miniature CMOS camera (Blackfly s-USB3, Flir) with a frame rate of 10 Hz using FlyCam2 software (Flir).

### Visual stimulation

For visual stimulation experiments, sinusoidal drifting gratings (2 Hz, 0.04 cycles/degree) with varied contrast were generated using custom-written functions based on Psychtoolbox in Matlab and presented on an LCD monitor at a distance of 20 cm from the right eye. Stimuli were presented for 2 seconds with a 5 second inter-stimulus interval.

### Data analysis

All analyses were conducted using custom-written scripts in MATLAB (Mathworks). SVM classifiers were trained using publicly available software ^50^.

#### Preprocessing of behavior data

Pupil diameter and facial movements were extracted from face videography using FaceMap ^12^. For subsequent analysis, facial movement is defined as the the first component of FaceMap-based decomposition. Singular value decomposition (SVD) was applied to the face movie to extract the principal components (PCs) explaining the distinct movements apparent on the mouse’s face. Wheel position was obtained from a linear angle detector attached to the wheel axle by unwrapping the temporal phase and then computing the traveled distance (cm). Locomotion speed was computed as the differential of the smoothed distance (cm/sec) using a 0.4 second window. Epochs of sustained locomotion and quiescence were extracted using change-point detection as described ^14^. High/low Pupil and Face epochs were extracted from within quiescence segments where z-score normalized values exceeded high/low thresholds of 60%/40% quantiles.

#### Preprocessing of mesoscopic imaging data

Imaging frames for green and red collection paths were grouped and down-sampled from 512×512 to 256×256 followed by an automatic ‘rigid’ transformation (imregtform, Matlab). In some cases, registration points were manually selected and a ‘similarity’ geometric transformation was applied. Detrending was applied using a low pass filter (*N* = 100, *f*_cutoff_ = 0.001*Hz*). Time traces were obtained using (Δ*F*/*F*)_*i*_ = (*F_i_* – *F_i,o_*)/*F_i,o_* where *F_i_* is the fluorescence of pixel *i* and *F_i,o_* is the corresponding low-pass filtered signal.

#### Hemodynamics correction

Hemodynamic artifacts were removed using a linear regression accounting for spatiotemporal dependencies between neighboring pixels ^14^. We used the approximate isosbestic excitation of GCaMP6 or GRAB_ACh3.0_ (395 nm) as a means of measuring activity-independent fluctuations in fluorescence associated with hemodynamic signals. Briefly, given two *p* × 1 random signals *y*_1_ and *y*_2_ corresponding to Δ*F/F* of *p* pixels for two excitation wavelengths “green” and “UV”, we consider the following linear model:

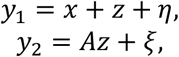

where *x* and *z* are mutually uncorrelated *p* × 1 random signals corresponding to *p* pixels of the neuronal and hemodynamic signals, respectively. *η* and *ξ* are white Gaussian *p* × 1 noise signals and *A* is an unknown *p* × *p* real invertible matrix. We estimate the neuronal signal as the optimal linear estimator for *x* (in the sense of Minimum Mean Squared Error):

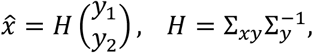

where 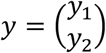 is given by stacking *y*_1_ on top of *y*_2_, 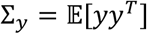 is the correlation matrix between *y* and 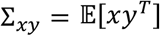 is the correlation matrix between *x* and *y*. The matrix ∑_*y*_ is estimated directly from the observations, and the matrix ∑_*xy*_ is estimated by^14^:

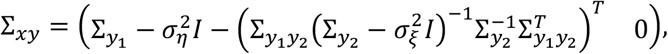

where 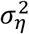 and 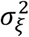 are the noise variances of *η* and *ξ*, respectively, and *I* is the *p* × *p* identity matrix. The noise variances 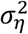 and 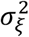 are evaluated according to the median of the singular values of the corresponding correlation matrices ∑_*y*1_ and ∑_*y*2_^51^. This analysis is usually performed in patches where the size of the patch, *p*, is determined by the amount of time samples available and estimated parameters. In the present study, we used a patch size of *p* = 9. The final activity traces were obtained by z-scoring the corrected Δ*F/F* signals per pixel.

#### Parcellation of mesoscopic data using LSSC

Functional parcellation of mesoscopic data was performed using Local Selective Spectral Clustering (LSSC)^28^. Briefly, this method identifies areas of co-activity by building a graph where nodes are pixels and edge weights are determined by pairwise similarities between activity traces of pixels obtained by the following kernel:

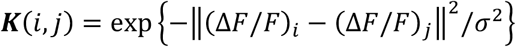

where σ is a parameter expressing a similarity radius. A row-stochastic matrix ***P*** is obtained by normalizing the rows such that ***P*** = ***D*^−1^*K***, where ***D***(*i, i*) = ∑_*j*_***K***(*i, j*). The matrix ***P*** can be viewed as a transition matrix of a Markov chain of the graph where ***P***(*i, j*) is the probability to jump from node (pixel) *i* to node (pixel) *j*. We obtain a non-linear embedding of pixels by calculating the *d* right eigenvectors with the largest eigenvalues of *P*:

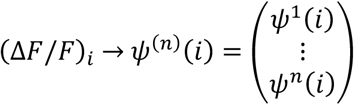

Overall, by taking *n* to be significantly smaller than the number of time samples, every pixel is represented by a lower dimensional embedding *ψ*^(*n*)^.

We evaluate the embedded representation *ψ*^(*n*)^ and calculate the spectral embedding norm ^52^ of every pixel *s_i_* = ||*ψ*^(*n*)^(*i*)||. LSSC uses an iterative approach for parcellating the brain where the inputs are the embedded representation of all pixels *ψ*^(*n*)^ and their corresponding norms, *s_i_*, *i* = 1, …, *p*, and lastly, a list of all pixels sorted by decreasing order of the embedding norm denoted by *l*. On each iteration the following operations are performed until coverage of at least *ϑ* percent of the mask brain pixels is assigned to parcels:

1. Select the first item on the list *l* (the pixel having the maximal norm, noted by *i*\*)
2. Select the axes in which *i*\* has the largest values, i.e., the subset: *L*_*i*\*_ = [*ℓ*_1_, ℓ_2_, …, *ℓ_d_i__* such that |*ψ^ℓ_1_^* (*i*)≥ *ϕ*^ℓ_2_^(*i*)|≥ |*ψ*^*ℓ*_3_^(*i*)|≥ ⋯
3. Obtain the pixels whose embeddings are closer to *ψ*^(*n*)^(*i*) than to the origin based on the axes *L*_*i*\*_ and assign them to the cluster *k*, i.e.:

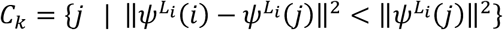
4. Remove the set *C_k_* from the list *l*: *l*←*l* \ *C_k_*
5. *k* ← *k* + 1
6. If at least *ϑ* percent of the mask of the brain is assigned to a specific parcel, then break.

The output is therefore a set of clusters {*C_k_*}where each clusters contains the pixels in that cluster. To increase robustness, we divided every session into 10 disjoint segments (folds), extracted the embedding on every fold and evaluated the embedding norm as the maximal value across all 10 folds. We refined the brain parcellation by merging parcels whose time traces are correlated more than a given threshold. Overlapping pixels were assigned to the parcel with closest centroid (in the embedding space). Additionally, unassigned isolated pixels (if any) were assigned to the (spatially) closest parcel. Isolated pixels within the borders of more than one parcel were assigned to the closest cluster (in the embedding space). Each animal and session was parcellated to reach a 95% coverage of the mask of the brain where clusters were merged based on a threshold of 0.99, resulting in <45 parcels per hemisphere. Time series for parcels were extracted by averaging values for all pixels within the parcel (see preprocessing of mesoscopic data above).

#### ROI extraction by LSSC

We also used LSSC to identify cell bodies from the two-photon imaging data. The overall approach is similar to the parcellation process except for the stopping condition, where iterations continue until a maximal number of cells is reached. In the refinement stage, identified cells that smaller than 15 pixels were discarded and overlapping regions were resolved by de-mixing ^28^.

#### Taylor expansion for estimating behavior as a function of neuronal activity

We formulate the link between temporal dynamics of neuronal activity 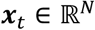 and an observed behavior *b_t_* as:

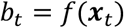

where *f* is an unknown function. Assuming that *f*(***x**_t_*) is 2 times differentiable, we can write its second-order Taylor’s expansion as:

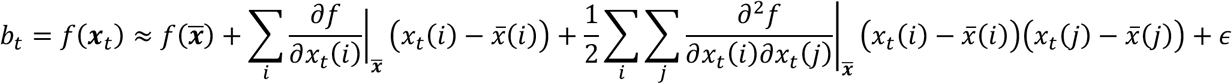

where 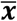 is the average neuronal activity (across time), and *ε* is the error of neglecting higher orders of ***x**_t_*. Simplifying this equation leads to:

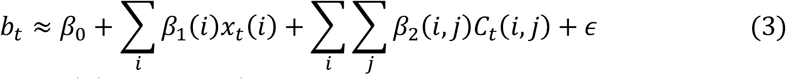

where 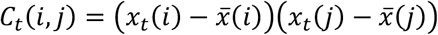 is the time trace of the instantaneous interaction between brain region *i* and brain region *j* and *β_n_, n* = 0,1,2 are the model parameters. Overall eq. (3) proposes a linear model for behavior based on two temporal signals - the activity *x_t_*(*i*) and the pairwise interaction *C_t_*(*i, j*), which is a nonlinear second-order function of elements of ***x**_t_*. Since the elements *x_t_*(*i*) and *C_t_*(*i, j*) are linearly independent for all *i, j* ∈ [1, *N*] can measure the decoding power of each of these two components ***x**_t_* and ***C**_t_* independently.

In eq. (3) the instantaneous interactions *C_t_*(*i, j*) are evaluated based on a single time point. In practice, estimating all pairwise interactions at a single point is highly sensitive to noise. Thus, we evaluate the interactions over a short sliding time window to obtain the sample covariance *Ĉ_t_*(*i, j*) as a smoothed and more robust estimation for the temporal evolution of *C_t_*(*i,j*):

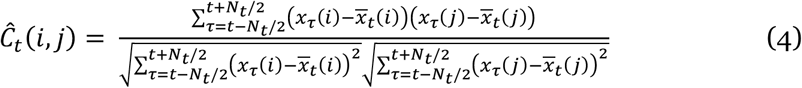

where 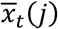 is the smoothed averaged activity:

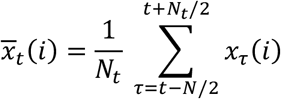

Inserting *Ĉ_t_*(*i,j*) into (3) leads to eqn. (1). Overall, 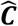 is a three-dimensional tensor of parcels by parcels by time, where each element *Ĉ_t_*(*i,j*) is a time trace of the instantaneous correlation coefficient between parcel *i* and parcel *j*. For most analyses, *N*_t_ was 30 (corresponding to a 3 second moving window). In all cases, the time-step was set to be 1 frame (0.1 second).

#### Riemannian projection of correlation matrices

Correlation matrices are Symmetric and Positive Definite (SPD, i.e., symmetric and full rank) and whose underlying geometry is a manifold shaped like a cone with a Riemannian metric (Supplemental Figure 2) ^53,54^. The distances between two correlation matrices on this cone is defined by the geodesic distance, the length of the arc connecting these matrices, whereas the Euclidean distance is not an accurate measure for this geodesic distance. To accurately capture distances between SPD matrices, Riemannian geometry is often used to project them onto a tangent Euclidean space where the geodesic length is approximated by the Euclidian distances between the corresponding projections. This evaluation becomes more accurate if the plane is tangent to the cone at a point that is relatively close to all relevant matrices, usually taken as their Riemannian mean.

Briefly, let {*C_k_*} be a set of *K* SPD matrices. Denote 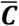 as the Riemannian mean of the set and 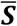 as its equivalent in the tangent plane. 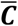 and 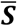 are calculated using the following iterative equations:

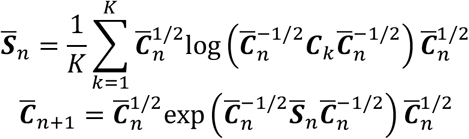

Where log (·) and exp (·) are the matrix logarithm and matrix exponential, respectively, and where the Euclidean mean is used to initialize: 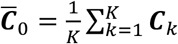. Convergence is obtained when the Frobenius norm of 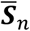 is smaller than a pre-set parameter 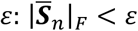.

The projections of {*C_k_*} onto the tangent plane to the cone at the Riemannian mean are given by:

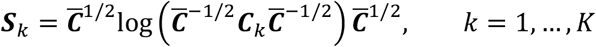

As presented previously ^55^, the pairwise distances, 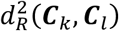 on the cone between correlation matrices {*C_k_*} can be approximated by the Euclidean distances between their corresponding projections {*S_k_*}:

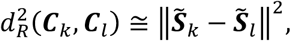

where 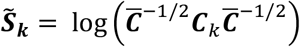. This method requires that all matrices {*C_k_*} would be full rank ^6,57^. In practice this is not always the case if the number of time points for evaluation of the correlation matrices is smaller than *p*, i.e. *N_T_* < *p*. Therefore, we add a regularization term, λ***I***, to each correlation matrix *C_k_*^58^ where *λ* is set to the median of the singular values of ***x**_t_* ^51^.

#### Dimensionality reduction by diffusion embedding

The series of matrices *C_t_* are symmetrical and therefore the dimension of their projections, {***S**_t_*}, is equal to 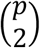 resulting in a high dimensional signal. To analyze the dynamics of this signal, we used diffusion geometry to obtain a low dimensional representation, capturing the dynamical properties of the correlation traces. Unlike LSSC where we reduce the dimension across time samples, in this case we reduce the dimension of parcels; we evaluated the *N_T_* × *N_T_* kernel matrix of {*S_t_*}:

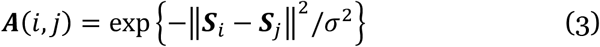

where *σ*, which is a scale parameter evaluated as the median of pairwise distances between each projected matrix and its *k*-nearest neighbors where *k*=20. Note that our results are highly robust to variation in this parameter (Supplemental Figure 2).

Normalizing the kernel ***A*** to be row-stochastic and taking the right eigenvectors leads to the low dimensional representation for the correlation traces:

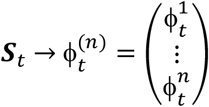

The size of the kernel matrix ***A*** is determined by the available time points recorded on each experiment (*N_T_* ≈10^4^ for all sessions). To reduce computational complexity involving eigenvalue decomposition of large matrices we used Nyström out-of-sample extension ^59^ as follows. We randomly choose a smaller subset of time points 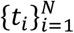, where *N* < *N_T_*. We obtain the normalized kernel using these time points and extract its eigen-decomposition *λ_k_*, 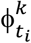, *k* = 1, …, *n*. We then extend this low dimensional representation to all time points:

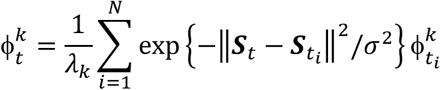

For comparison to using Riemannian geometry for calculating distances in the kernel, we carried out similar diffusion embedding based on Euclidean distances (Supplemental Figure S2).

#### Dimensionality reduction by principal component analysis

As a comparison to LSSC, we also used principal component analysis to reduce the dimensionality of widefield data (Supplemental Figure S2). Principal components were derived using the ‘pca’ function in Matlab.

#### Visual response analysis

Visual responses were evaluated, per parcel, as the difference between peak response during stimulus presentation and the average activity during the preceding two seconds. The responses were averaged per contrast value and normalized by the response to 100% contrast. To quantify the accuracy with which visual responses are encoded by visual activity, or embedded network activity/correlations, we trained a binary classifier (linear SVM, libsvm) to separate the visual response and the two seconds prior to stimulus onset. We used 10-fold cross validation to estimate the classification accuracy for every contrast value based on each predictor.

#### Modeling Behavior

Behavioral variables (pupil, facial movements, running speed) were modeled using linear ridge regression with 10-fold cross validation. Each session was divided into 10-disjoint continuous segments, where on each fold one segment was set aside for testing and the other segments were used for training. We assessed the predictive power of neuronal activity using the following predictors: raw activity and smoothed activity (using a 3 second moving window). For pairwise correlations, we used: raw correlation traces, diffusion embedding of correlation traces using either Euclidean or Riemannian distances. To directly compare the predictive power of activity versus embedded correlations, we combined these predictors and evaluated the goodness of fit of the joint model. We then shuffled either activity or embedded correlations through time and trained the resulting model to assess 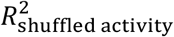 and 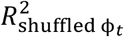 *ϕ_t_*.

#### Reconstruction Error

Reconstruction error of diffusion embedding of functional connectivity was evaluated by:

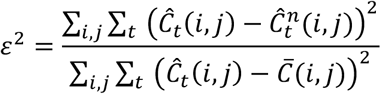

where 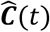 is the pairwise functional connectivity of brain parcels, 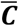 is the temporal average and where 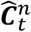 is the reconstruction of 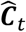 based on *n* leading components of its embedded trace 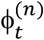:

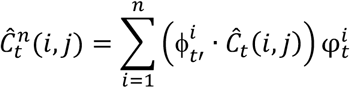

where 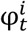 are the left eigenvectors of the normalized kernel matrix.

#### Modeling correlations data by embedding

To quantify the relationship between the embedding of functional connectivity across the cortex ϕ_*Z*_ and the time-varying correlation between specific pairs of parcels, we used linear regression (10-fold cross validation) and obtained an R^2^ value for every pair-wise correlation trace. To match LSSC parcels across animals, we identified the LSSC parcels whose center of mass were closest to each Allen Atlas brain parcel (23 parcels overall in a single hemisphere) and extracted a 23 × 23 matrix of R^2^ values per session. We averaged these matrices across animals and extracted the rows corresponding to individual parcels. Each row was then represented as a separate brain map image, color-coded by the R^2^ value corresponding to the correlation between the target (specific to that image) and each of the other parcels.

#### Evaluating Integrated Network Configuration

We evaluated a time trace of the average participation coefficient (across parcels), based on the correlation time traces (3 seconds) and an arbitrary partition of the brain using a line bisecting the neocortex. The participation coefficient is calculated per parcel in a given time window *t* as:

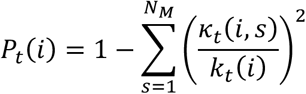

where *κ_t_*(*i, s*) is the sum of the correlations (absolute values) between parcel i and all the parcels in sub-module s and *k_t_*(*i*) is the sum of all correlations (absolute values) between parcel i and all other parcels. Therefore, a segregated network would be characterized by high connectivity between parcels related to the same module and low connectivity between sub-modules. In this case the average participation coefficient (across parcels) would approach zero. The opposite would happen in an integrated network where parcels communicate outside their sub-modules just as much as they do within their sub-modules. In this case the average participation coefficient would approach 1.

We then evaluated the difference of the behavior variables (pupil size, facial movement, and locomotion speed) between time points where the network was in an extreme integrated state (top 10% participation coefficient) and extreme segregated state (bottom 10% participation coefficient) ^35^. By rotating the line in 30, 60, 90, 120, 150 degrees we measured the delta of behavior variables on integrated and segregated configurations based on different ways for partition of the brain into sub-networks.

#### Instantaneous Multimodal Connectivity

Denote 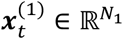 as the neuronal activity of *N*_1_ brain parcels at time *t* and denote 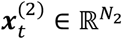 as the neuronal activity of *N*_2_ cells at time *t*. We define ∑_*t*_(*i, j*) as a 3-dimensional tensor of *N*_1_ over *N*_2_ over time, expressing the dynamics of multimodal connectivity between cells and parcels:

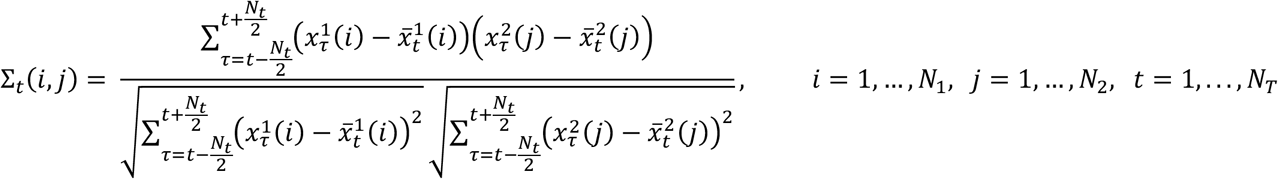

where *N_T_* is the numer of time points. For a given time point *t*, the matrix ∑_*t*_, is non-symmetric and therefore not bound to the Riemannian cone (as opposed to the sample covariance matrices 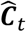, which are SPD). Therefore, we used Euclidean distances to evaluate the diffusion kernel between correlation matrices related to different time points:

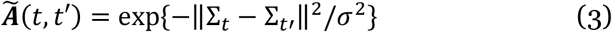

where *σ* is a scale parameter evaluated as the median of pairwise distances between each correlation matrix and its *k*-nearest neighbors, where *k*=200. Note that as for embedding of correlation matrices, our results here are also highly robust to variation in *k* (Supplemental Figure 6). Here we again reduce computational complexity using Nyström extension and evaluate the kernel 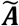 based on a smaller randomly selected sub-set of time points, *t*′ = 1,.., *N*, where *N* < *N_T_*. We normalize the kernel matrix to be row-stochastic and obtain its eigen-decomposition:

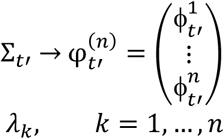

We then extend this representation to all time points *t* = 1,.., *N_T_*:

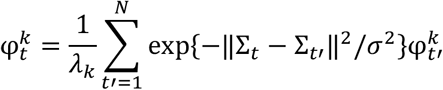

#### Standard Deviation of Multimodal Connectivity

We estimate the average variability of correlations between a given cell and the mesoscopic cortical network as:

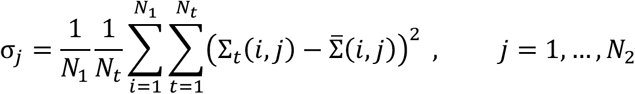

where 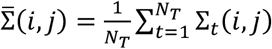.

## Supplemental Figures

**Supplemental Figure 1.**
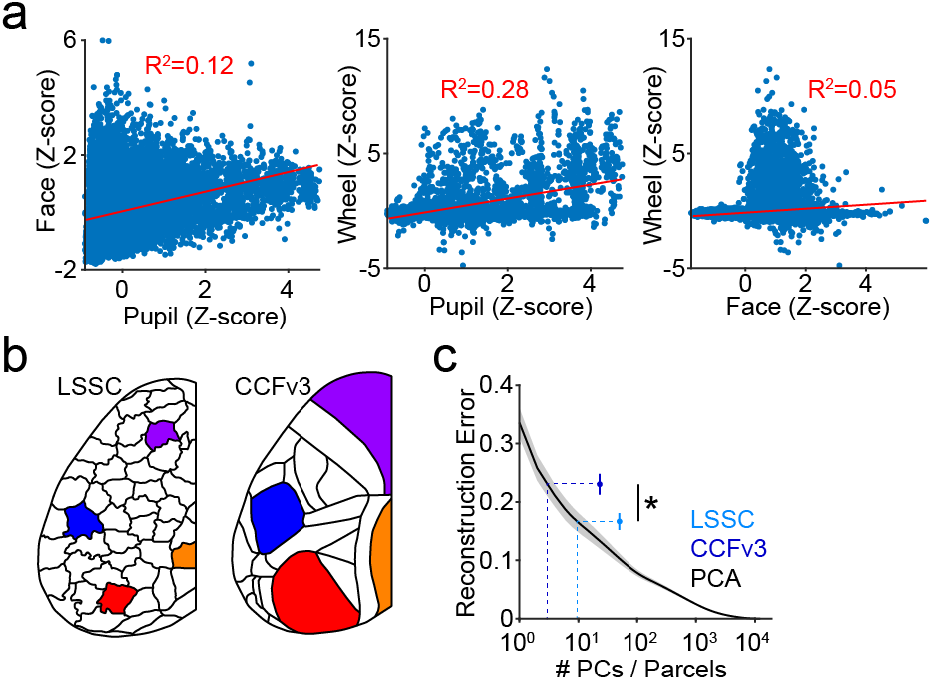
Comparison of parcellation methods and average neural signaling across behavioral state. **a**, Example scatter plots illustrating the relationship between Z-scored facial movement (Face), Pupil diameter (Pupil), and Locomotion (Wheel) for a representative subject. Pearson’s correlation line and goodness of fit (R^2^) are shown in red. **b**, Example LSSC-based functional parcellation (left) and CCFv3-based anatomical parcellation (right) for the data in Figure 1. Coloring indicates approximately matching areas (MO_s_ purple, SS_P_ blue, VIS_P_ red, RS orange). **c**, Reconstruction error for raw activity using either LSSC- or CCFv3-based parcellation or principal component analysis (PCA). PCA indicates error versus number of components included in the model. LSSC and CCFv3 points indicate the number of parcels obtained for each method.

**Supplemental Figure 2.**
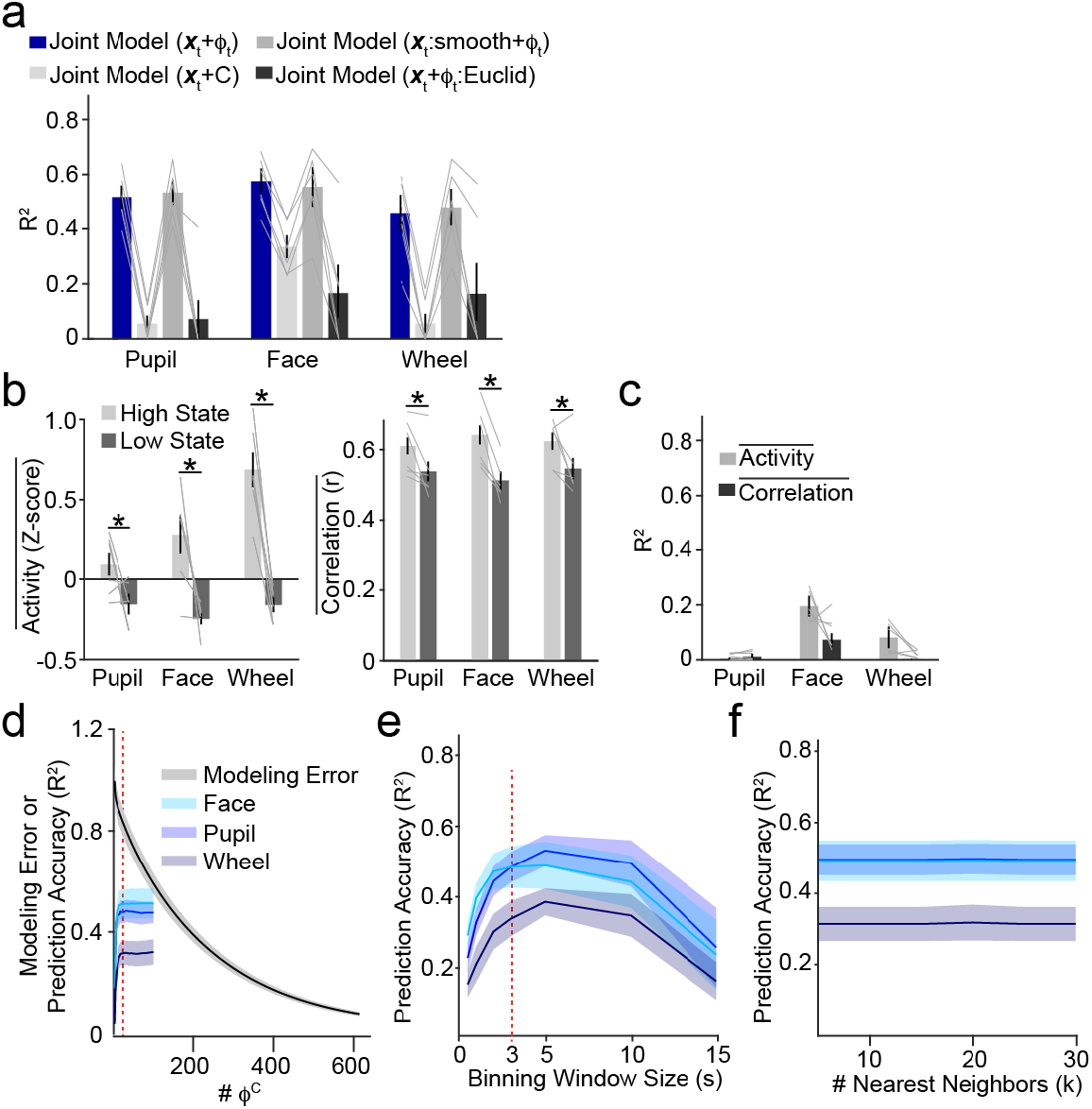
Comparison of behavioral prediction across different models. **a**, Widefield imaging population data of showing average (±SEM) prediction accuracy (R^2^) for modeling behavior variables using a joint model of activity and embedded correlations (replicated from Figure 2, blue), joint model with raw correlations (light gray), joint model with smoothed (windowed) activity (medium gray), and joint model using Euclidean embedding (dark gray). **b**, Population data showing average (± SEM) of cortical activity (left) and correlations (right) across all LSSC-based parcels, comparing low versus high divisions of the indicated behavioral state. * indicates p<0.05 (see Main Text). **c**, Prediction accuracy for modeling behavioral variables using the average activity or correlations across all cortical parcels. **d**, Modeling reconstruction error of embedded correlations (black) or prediction error of behavioral metrics (shades of blue) versus the number of *ϕ_c_* components. Shaded areas indicate ±SEM. **e**, Prediction accuracy (R^2^) for behavioral metrics using 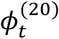 versus sliding temporal window size for calculating correlations. **f**, Prediction accuracy (R^2^) of behavioral metrics using 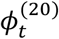 versus the number of neighbors used for evaluating the scale factor of the diffusion kernel. * indicates p<0.05.

**Supplemental Figure 3.**
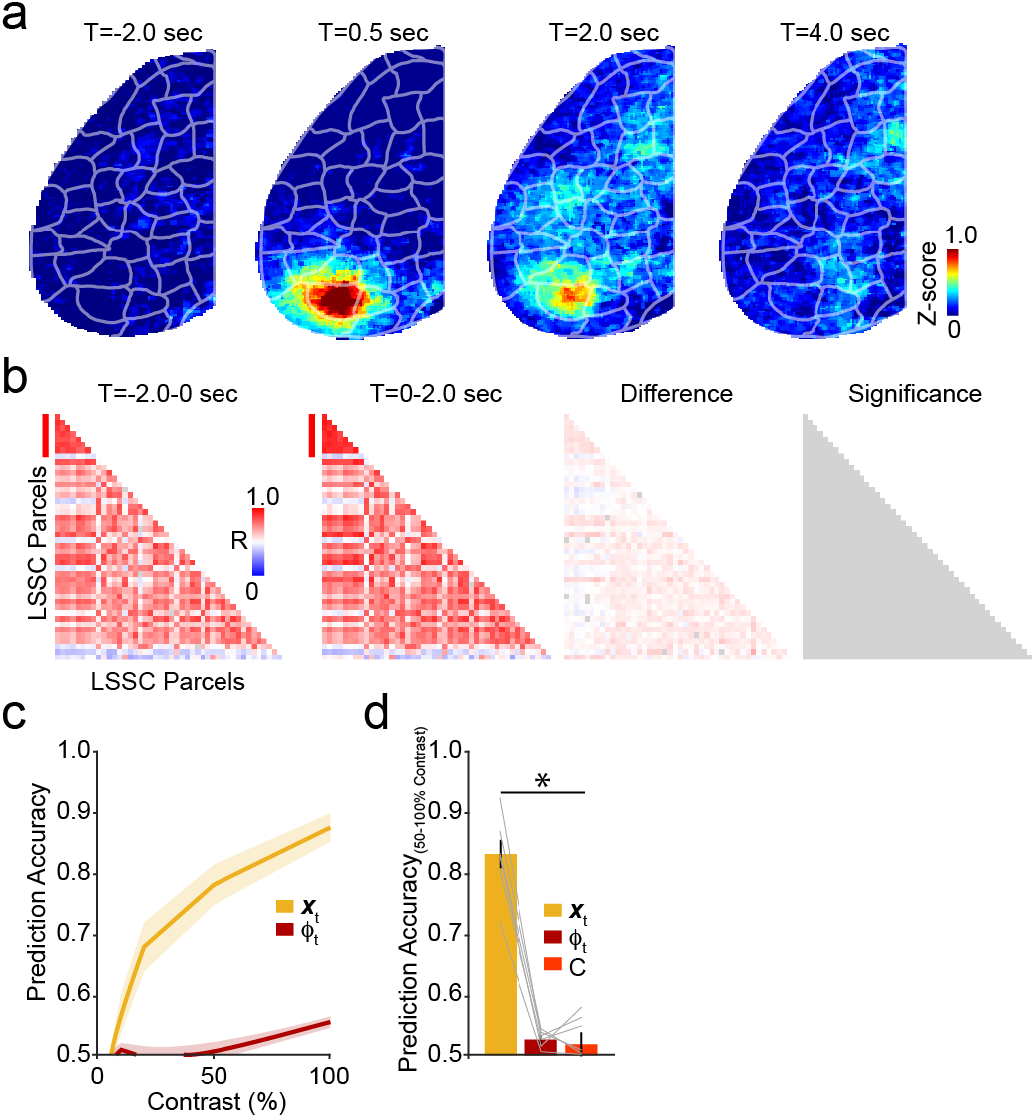
Network representation of visually-evoked activity. **a**, Example average (n=50 stimuli, one animal) mesoscopic imaging frames showing evoked visual responses (100% contrast) relative to stimulus onset. **b**, Example average correlation matrices corresponding to data from (a) showing functional connectivity of parcels before (first panel) and after (second panel) visual stimulation (100% contrast). The differences in correlation for each parcel (third panel) and the significant parcels (fourth panel) are shown. All cells are gray, indicating no significant differences (p<0.05, permutation test, Benjamini-Hotchberg multiple comparisons correction) for any parcel pair. **c**, Prediction accuracy for detecting the presentation of a visual stimulus for varying contrasts using activity in visual cortex (yellow) or *ϕ_t_* (dark red). Shaded areas indicate SEM (n=6 mice). **d**, Population averages (n=6 mice) showing prediction accuracy of visual responses (averaging across trials with ≥50% stimulus contrast) for visual cortex activity (yellow), *ϕ_t_* (red), and raw correlations (red). * indicates p<0.05 (ANOVA, Tukey’s post-hoc tests, see main text).

**Supplemental Figure 4.**
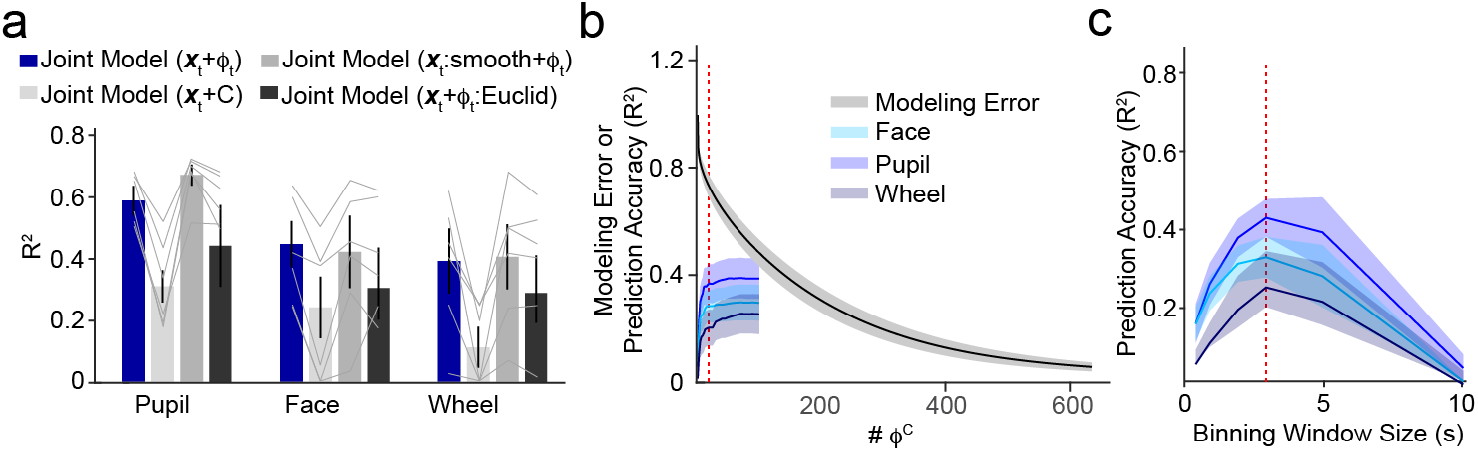
Diffusion embedding of cellular correlations is robust to choice of modeling parameters. **a**, Cellular imaging population data of showing average (±SEM) prediction accuracy (R^2^) for modeling behavior variables using a joint model of activity and embedded correlations (replicated from Figure 3, blue), joint model with raw correlations (light gray), joint model with smoothed (windowed) activity (medium gray), and joint model using Euclidean embedding (dark gray). **b**, Modeling reconstruction error of correlations (black) or prediction error of behavioral metrics (shades of blue) versus the number of *ϕ_c_* components. Shaded areas indicate ±SEM. **c**, Prediction accuracy (R^2^) of behavioral metrics using 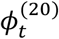 versus sliding temporal window size for calculating correlations.

**Supplemental Figure 5.**
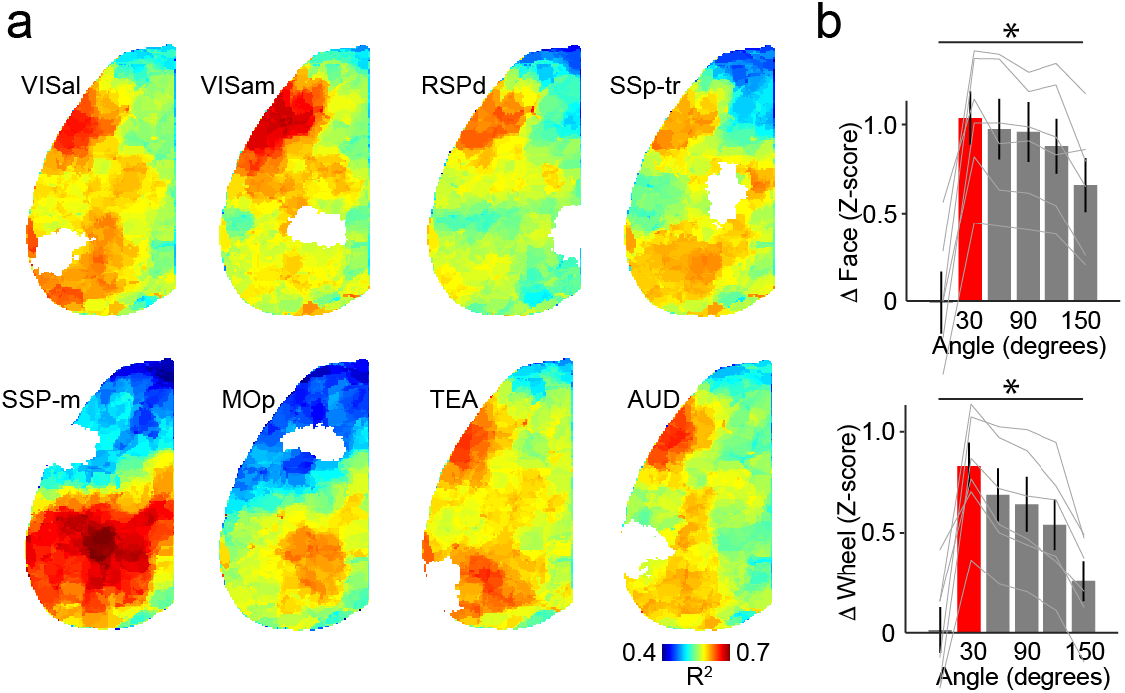
Dynamic connectivity reveals consistent spatial organization of cortical subnetworks. **a**, Average (n=6 mice) maps showing mean R^2^ values for modeling the pairwise correlations of each cortical parcel with the indicated target parcel (showing in white). **b**, Population data showing the average (±SEM) difference in facial movement (top) and wheel speed (bottom) for epochs corresponding to maximal and minimal network segregation versus a line angle bisecting LSSC parcels (as in Figure 4). * indicates p<0.05 (ANOVA).

**Supplemental Figure 6.**
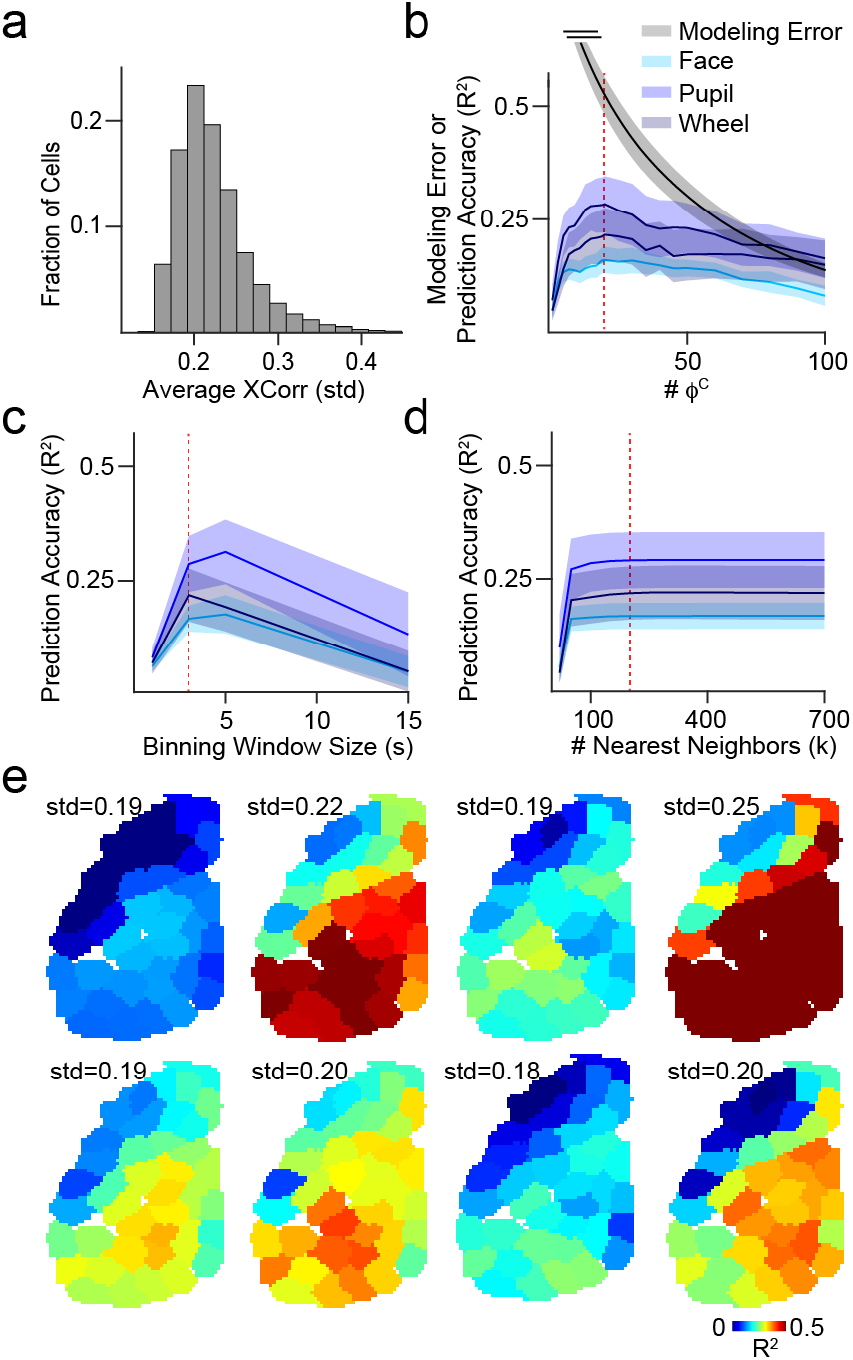
Diffusion embedding of dual mesoscopic and cellular correlations is robust to choice of modeling parameters. **a**, Population data for the standard deviation of multimodal correlation values over time, for each cell averaged across all parcels. **b**, Modeling reconstruction error of correlations (black) or prediction error of behavioral metrics (shades of blue) versus the number of *ϕ_c_* components. Shaded areas indicate ±SEM. **c**, Prediction accuracy (R^2^) of behavioral metrics using 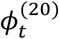 versus sliding temporal window size for calculating correlations. **d,** Prediction accuracy (R^2^) of behavioral metrics versus the number of neighbors used for evaluating the scale factor of the diffusion kernel. **e,** Additional example maps for different cells showing R^2^ values for modeling the correlation of the cell with each cortical parcel using the overall diffusion embedding.

